# Protein Phosphatase 1 regulates atypical mitotic and meiotic division in *Plasmodium* sexual stages

**DOI:** 10.1101/2021.01.15.426883

**Authors:** Mohammad Zeeshan, Rajan Pandey, Amit Kumar Subudhi, David J. P. Ferguson, Gursimran Kaur, Ravish Rashpa, Raushan Nugmanova, Declan Brady, Andrew R. Bottrill, Sue Vaughan, Mathieu Brochet, Mathieu Bollen, Arnab Pain, Anthony A. Holder, David S. Guttery, Rita Tewari

## Abstract

PP1 is a conserved eukaryotic serine/threonine phosphatase that regulates many aspects of mitosis and meiosis, often working in concert with other phosphatases, such as CDC14 and CDC25. The proliferative stages of the malaria parasite life cycle include sexual development within the mosquito vector, with male gamete formation characterized by an atypical rapid mitosis, consisting of three rounds of DNA synthesis, successive spindle formation with clustered kinetochores, and a meiotic stage during zygote to ookinete development following fertilization. It is unclear how PP1 is involved in these unusual processes. Using real-time live-cell and ultrastructural imaging, conditional gene knockdown, RNA-seq and proteomic approaches, we show that *Plasmodium* PP1 is implicated in both mitotic exit and, potentially, establishing cell polarity during zygote development in the mosquito midgut, suggesting that small molecule inhibitors of PP1 should be explored for blocking parasite transmission.

## Introduction

Cell cycle progression involves sequential and highly ordered DNA replication and chromosome segregation in eukaryotes^1, 2^, which is tightly controlled and coordinated by reversible protein phosphorylation catalysed by protein kinases (PKs) and protein phosphatases (PPs)^3^. Numerous studies have highlighted the importance of the phosphoprotein phosphatase (PPP) family in regulating mitosis in model eukaryotic organisms, often working in conjunction with CDC25 and CDC14 phosphatases, as key regulators of mitotic entry and exit, respectively^4,5,6,7^.

Protein Phosphatase 1 (PP1) is a member of the PPP family and is expressed in all eukaryotic cells. It plays a key role in the progression of mitosis through dephosphorylation of a large variety of proteins, including mitotic kinases (such as cyclin-dependent kinase 1 (CDK1)^8, 9, 10, 11^, and regulators of chromosome segregation^12^. Chromosome segregation is orchestrated at the kinetochore, a protein complex that assembles on the centromeres, located at the constriction point of sister chromatids to facilitate and monitor attachment of the sister chromatids to spindle microtubules^13, 14^. Correct attachment of the kinetochore to spindle microtubules is regulated by the KMN (KNL1, MIS12 and NDC80) protein network^15^, which in mammalian cells integrates the activities of at least five protein kinases (including MPS1, Aurora B, BUB1, PLK1, and CDK1) and two protein phosphatases (PP1 and PP2A-B56). The KMN network mediates kinetochore-microtubule attachments and scaffolds the spindle assembly checkpoint (SAC) to prevent chromosome segregation until all sister chromatids are properly connected to the spindle^16^. The orchestration of reversible protein phosphorylation is crucial to control the spatial-temporal progression of the cell cycle and PP1 has a key role in this process, in particular during mitotic exit. Throughout mitosis, PP1 is inhibited by the cyclin-dependent kinase (CDK1)/cyclin B complex and Inhibitor 1^17, 18^; however, during mitotic exit concomitant destruction of cyclin B and reduced activity of CDK1 through dephosphorylation by CDC14 results in subsequent reactivation of PP1 via autophosphorylation and completion of mitotic exit^10, 17, 19,20, 21^.

In *Plasmodium*, the causative organism of malaria, there is a single PP1 orthologue, which is expressed throughout the parasite’s complex life-cycle^22^ and located in both nucleus and cytoplasm^23^. *P. falciparum* PP1 (*Pf*PP1) shares 80% identity with human PP1c, and likely has a conserved tertiary and secondary structure containing 9 α-helices and 11 β-strands^24^. However, *Pf*PP1 lacks part of the 18 amino-acid motif at the C-terminus of human PP1, which contains a threonine residue (Thr320) that is phosphorylated by CDC2 kinase^25^ and dynamically regulates entry into mitosis^24^. Previous studies determined that *P. falciparum* PP1 functionally complements the *Saccharomyces cerevisiae* glc7 (PP1) homologue^26^, and subsequent phylogenetic analyses revealed that homologues of the phosphatases CDC14 and CDC25 are absent from *Plasmodium*^23^. Genetic screens and inhibitor studies have shown that PP1 is essential for asexual blood stage development^23, 27, 28, 29^ (also reviewed^24^), in particular for parasite egress from the host erythrocyte^30^. Most studies in *Plasmodium* have been focused on these asexual blood stages of parasite proliferation, and very little is known about the importance of PP1 for transmission stages within the mosquito vector. Recent phosphoproteomic studies and chemical genetics analysis have identified a number of potential *Plasmodium falciparum* (Pf) PP1 substrates modified during egress, including a HECT E3 protein-ubiquitin ligase and GCα, a guanylyl cyclase with a phospholipid transporter domain^30^. Other biochemical studies in *Plasmodium* have identified numerous PP1-interacting partners (PIPs) that are structurally conserved and regulate PP1 activity, including LRR1 (a human SDS22 orthologue), Inhibitor 2 and Inhibitor 3. Other PIPs including eif2β, and GEXP15 have also been identified^9, 24, 31, 32, 33^. Synthetic peptides containing the RVxF motif of Inhibitor 2 and Inhibitor 3, and the LRR and the LRR cap motif of *Pf*LRR1 significantly reduce *P. falciparum* growth in vitro^34, 35^ and regulate *Pf*PP1 phosphatase activity^35, 36, 37^. However, little is known regarding how PP1 is involved in regulating mitosis and meiosis in the absence of CDC14 and CDC25.

The *Plasmodium* life-cycle is characterised by several mitotic stages and a single meiotic stage (reviewed in^38, 39, 40^). However, in this organism kinetochore dynamics and chromosome segregation are poorly understood, especially in mitosis during male gametogenesis. This is a very rapid process with three rounds of spindle formation and genome replication from 1N to 8N within 12 minutes. In addition, little is known about the first stage of meiosis that occurs during the zygote to ookinete transition. We have followed recently the spatio-temporal dynamics of NDC80 throughout mitosis in schizogony, sporogony and male gametogenesis, and during meiosis in ookinete development^41^, using approaches that offer the opportunity to study the key molecular players in these crucial stages of the life cycle. Although PP1 has been partially characterised and shown to have an essential role during asexual blood stage development in the vertebrate host^23^, the role and importance of PP1 during sexual stage development in the mosquito is completely unknown.

Here, using the *Plasmodium berghei* (Pb) mouse model of malaria, we determined the importance of PP1 during the sexual stages within the mosquito vector. PP1 expression and location were studied using the endogenous, GFP-tagged protein and co-localisation with the kinetochore marker, NDC80, to follow progression through chromosome segregation during male gamete formation and zygote differentiation. Using a conditional gene knockdown approach we examined how PP1 orchestrates atypical mitosis and meiosis, and investigated the ultrastructural consequences of PP1 gene knockdown for cell morphology, nuclear pole multiplication and flagella formation during male gamete formation. RNA-Seq analysis was used to determine the consequence of PP1 gene knockdown on global transcription, which disclosed a marked differential expression of genes involved in reversible phosphorylation, motor activity and the regulation of cell polarity. Proteomics studies identified motor protein kinesins as interacting partners of PP1 in the gametocyte. The knockdown of PP1 gene expression blocks parasite transmission by the mosquito, showing that this protein has a crucial function in *Plasmodium* sexual development during both mitosis in male gamete formation and meiosis during zygote to ookinete differentiation.

## Results

### PP1-GFP has a diffuse subcellular location during asexual blood stage schizogony and forms discrete foci during endomitosis

To examine the spatio-temporal expression of *Plasmodium* PP1 in real-time during cell division, we generated a transgenic *P. berghei* line expressing endogenous PP1 with a C-terminal GFP tag. An in-frame *gfp* coding sequence was inserted at the 3’ end of the endogenous *pp1* locus using single crossover homologous recombination (**Fig S1A**), and successful insertion was confirmed by diagnostic PCR (**Fig S1B**). Western blot analysis of schizont protein extracts using an anti-GFP antibody revealed a major 62-kDa band, the expected size for PP1-GFP protein, compared to the 29 kDa WT-GFP (**Fig S1C**).

During asexual blood stage proliferation, schizogony is characterised by multiple rounds of asynchronous nuclear division without chromosome condensation or cytokinesis. Nuclear division is a closed mitotic process without dissolution-reformation of the nuclear envelope and with the spindle-pole body (SPB)/microtubule-organising centre (MTOC) embedded within the nuclear membrane^42^. It results in a multinucleated coenocyte termed a schizont, which is resolved by cytokinesis at the end of schizogony into individual merozoites. Live cell imaging of *P. berghei* asexual blood stages revealed a diffuse cytoplasmic and nuclear distribution of PP1-GFP, together with a distinct single focus of concentrated PP1-GFP in the nucleus of early trophozoite stage (**Fig 1A**), representing the early S-phase of the cell cycle when DNA synthesis starts. As schizogony proceeds, the diffuse distribution of PP1-GFP remained; however, in early schizonts each cell displayed two distinct PP1-GFP foci in close association with the stained nuclear DNA. These pairs of PP1-GFP foci became clearer during the middle and late schizont stages but following merozoite maturation and egress the intensity of the foci diminished (**Fig 1A**).

**Fig. 1.**
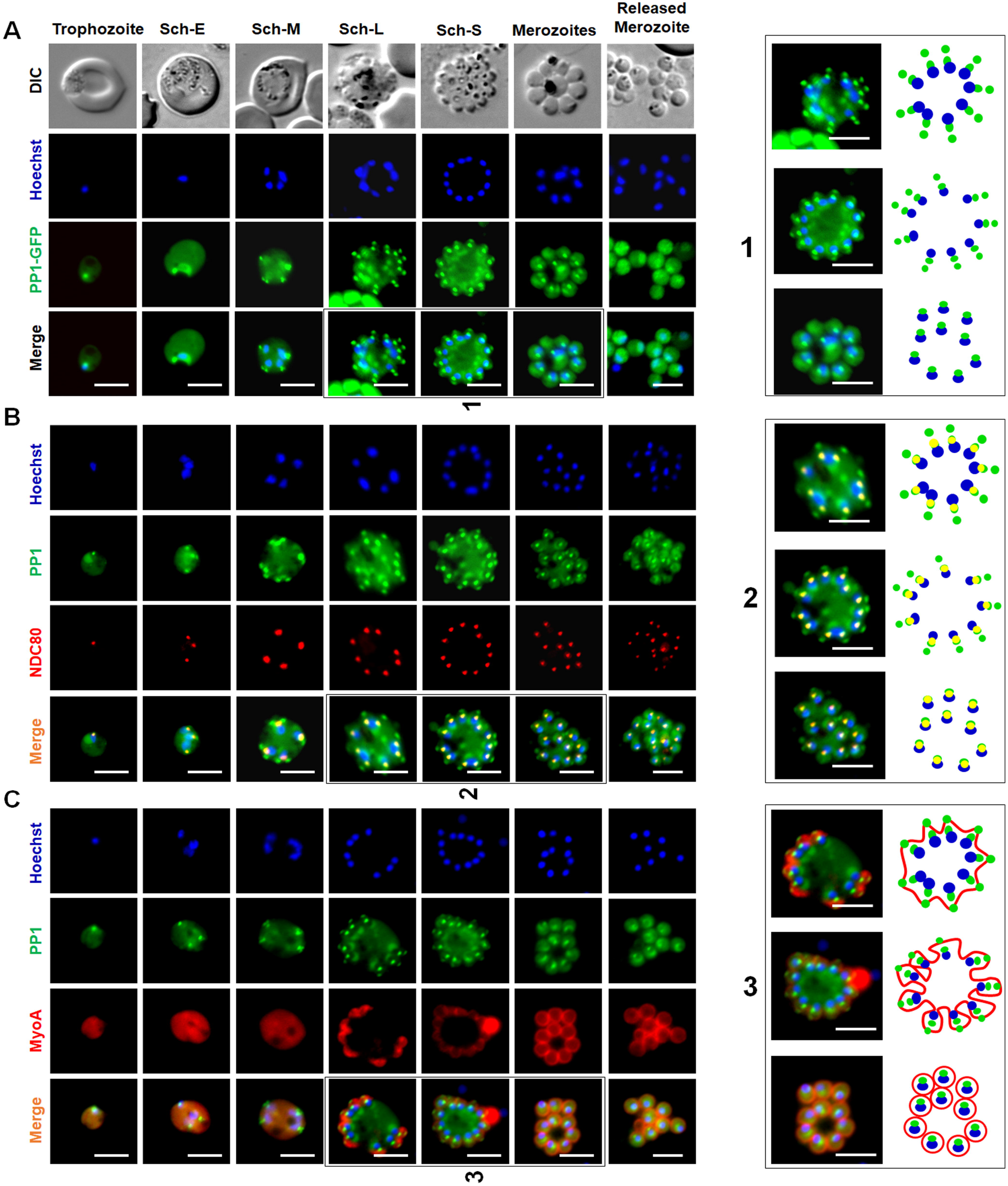
Location of PP1 during asexual blood stage schizogony and its association with kinetochore (Ndc80) and glideosome (MyosinA) **(A)** Live cell imaging of PP1-GFP (Green) showing its location during different stages of intraerythrocytic development and in free merozoites. DIC: Differential interference contrast; Hoechst: stained DNA (blue); Merge: green and blue images merged. A schematic guide showing the locations of PP1GFP foci during segmentation of merozoites is depicted in right hand panel. **(B)** Live cell imaging showing location of PP1–GFP (green) in relation to the kinetochore marker NDC80-mCherry (red) and DNA (Hoechst, blue). Merge: green, red and blue images merged. A schematic guide showing the locations of PP1GFP foci with NDC80-mCherry during segmentation of merozoites is depicted in right hand panel. **(C)** Live imaging showing the location of PP1–GFP (green) in relation to inner membrane complex marker MyoA-mCherry (red) and DNA (Hoechst, blue) during different stages of intraerythrocytic development and in extracellular merozoites. A schematic guide showing the locations of PP1GFP foci with NDC80-mCherry and MyoA-mCherry during segmentation of merozoites is depicted in the three right-hand panels. Merge: green, red and blue images merged. Sch-E (Early schizont), Sch-M (Middle schizont), Sch-L (Late schizont) Sch-S (Segmented schizont). In all panels, scale bar = 5 µm.

To study further the location of the PP1-GFP foci throughout mitosis, we generated by genetic cross parasite lines expressing either PP1-GFP and the kinetochore marker NDC80-mCherry^41^ or PP1-GFP and the inner membrane complex (IMC)-associated myosin A (MyoA)-mCherry. Live cell imaging of these lines revealed co-localisation of PP1-GFP and NDC80-mCherry foci close to the DNA of the nucleus through blood stage development, and especially during late schizogony and segmentation (**Fig 1B**); whereas the PP1-GFP foci at the outer periphery of the cell showed a partial co-location with MyoA-mCherry, and only during mid- to late schizogony (**Fig 1C**). Thus, in schizonts there is a concentration of PP1-GFP at the kinetochore and at a transient peripheral location during specific stages of schizogony, as well as the diffuse distribution throughout the cytoplasm (**Fig.1**).

### PP1-GFP is enriched on kinetochores during chromosome segregation associated with putative mitotic exit in male gametogony

To study PP1 expression and location through the three rounds of DNA replication and chromosome segregation prior to nuclear division during male gametogony, we examined PP1-GFP by live-cell imaging over a 15-minute period following male gametocyte activation. Following activation with xanthurenic acid and decreased temperature, male gametocytes undergo three rounds of DNA replication and mitotic division followed by chromosome condensation and exflagellation, resulting in eight gametes^43^. Before activation (0 min), PP1-GFP was detected with a diffuse location throughout gametocytes (**Fig 2A**). At one minute post-activation, PP1-GFP accumulated at one end of the nucleus at a single focal point (**Fig 2A**). After 2-3 min two distinct foci were observed on one side of the nucleus, concurrent with the first round of chromosome replication/segregation (**Fig 2A**). Subsequently, four and eight PP1-GFP foci were observed at 6 to 8 min and 8 to 12 min post-activation, respectively, corresponding to the second and third rounds of chromosome replication/segregation. These discrete PP1-GFP foci dispersed prior to karyokinesis and exflagellation of the mature male gamete 15 min post-activation, leaving residual protein remaining diffusely distributed throughout the remnant gametocyte and flagellum (**Fig 2A**).

**Fig. 2.**
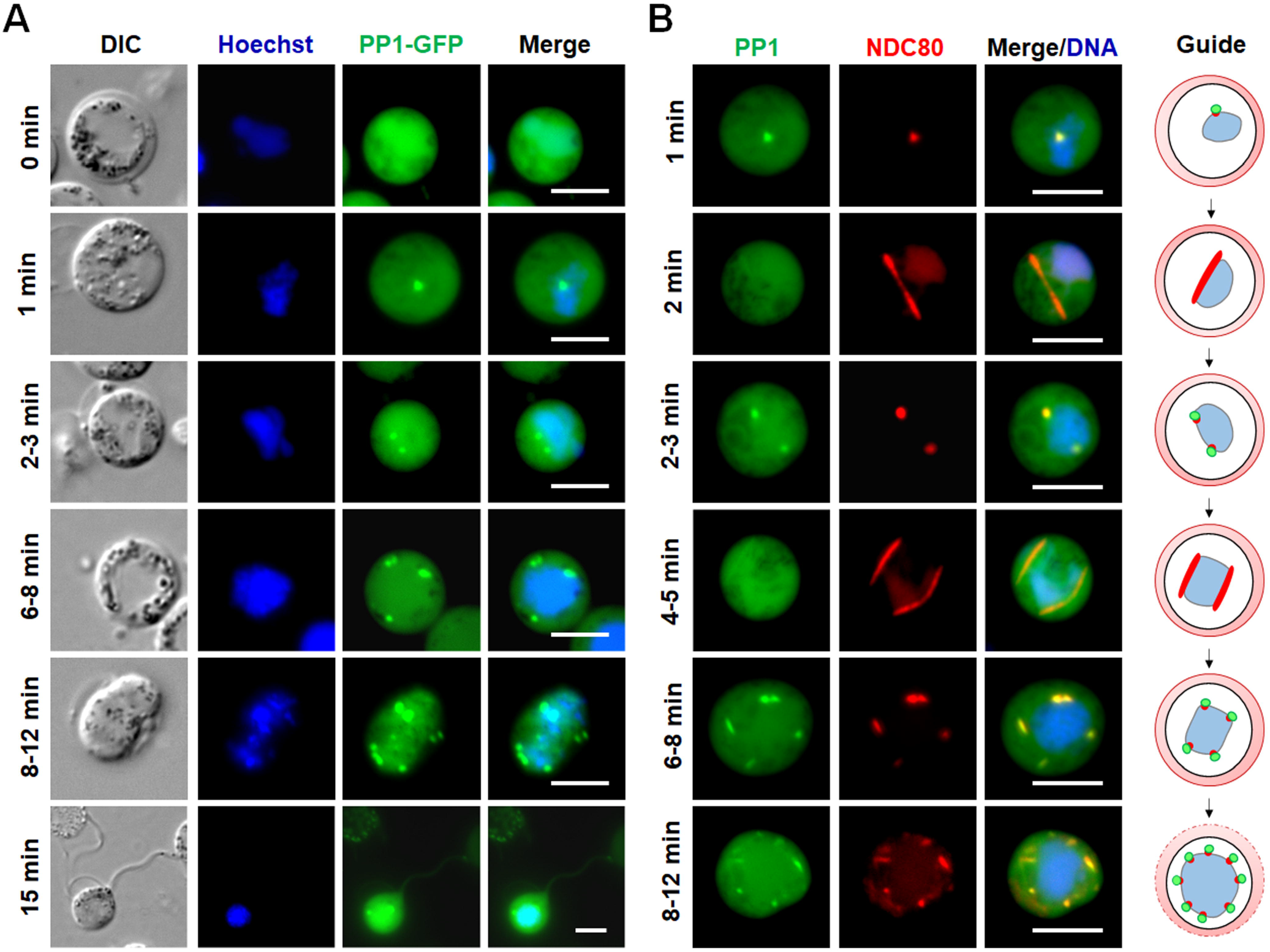
The location of PP1 and its association with the kinetochore during chromosome segregation in male gametogony. **(A)** Live-cell imaging of PP1-GFP during male gametogony showing an initial diffused localization before activation and focal points after activation in the later stages (shown as minutes post activation). Panels are DIC, Hoechst (blue, DNA), PP1-GFP (green) and Merge (green and blue channels). **(B)** Live-cell imaging of parasite line expressing both PP1-GFP and NDC80-mCherry showing location of PP1 (green) and NDC80 (red) in male gametocytes at different time points after activation. A schematic guide showing the locations of PP1GFP foci with DNA and NDC80-mCherry during male gametogony is depicted in the right panel. Merge/DNA is green, red and blue (Hoechst, DNA) channels. In both panels, scale bar = 5 µm.

To investigate further the location of PP1-GFP during spindle formation and chromosome segregation, the parasite line expressing both PP1-GFP and NDC80-mCherry was examined by live cell imaging to establish the spatio-temporal relationship of the two proteins. We found that the discrete PP1-GFP foci colocalized with NDC80-mCherry at different stages of male gametogony, including when up to eight kinetochores were visible (**Fig 2B**), but PP1-GFP was not associated with the arc-like bridges of NDC80-mCherry representing the spindle^41^ at 2 min and 4 to 5 min post activation, suggesting that PP1-GFP is only increased at the kinetochore during initiation and termination of spindle division (**Fig 2B**).

### PP1-GFP may determine apical polarity during zygote-ookinete development and has a nuclear location on kinetochores during meiosis

After fertilisation in the mosquito midgut, the diploid zygote differentiates into an ookinete within which the first stage of meiosis occurs. During this process DNA is duplicated to produce a tetraploid cell with four haploid genomes within a single nucleus in the mature ookinete. PP1 is known to be crucial during meiotic chromosome segregation^12^, and therefore we analysed the spatio-temporal expression of PP1-GFP during ookinete development using live cell imaging. PP1-GFP was expressed in both male and female gametes with a diffuse distribution, along with a single focus of intense fluorescence at one end (potentially the basal body) of each male gamete (**Fig. 3A**). Initially, the zygote also had a diffuse PP1-GFP distribution, but after two hours (stage I) an enriched focus developed at the periphery of the zygote, marking the point that subsequently protruded out from the cell body and developed into the apical end of the ookinete. A strong PP1-GFP fluorescence signal remained at this apical end throughout ookinete differentiation (**Fig 3A**). In addition, PP1-GFP was enriched in the nucleus at four discrete foci in mature ookinetes (**Fig 3A**). Analysis of the parasite line expressing both PP1-GFP and NDC80-mCherry showed that these four foci correspond to the kinetochores of meiotic development in the ookinete (**Fig 3B**).

**Fig. 3.**
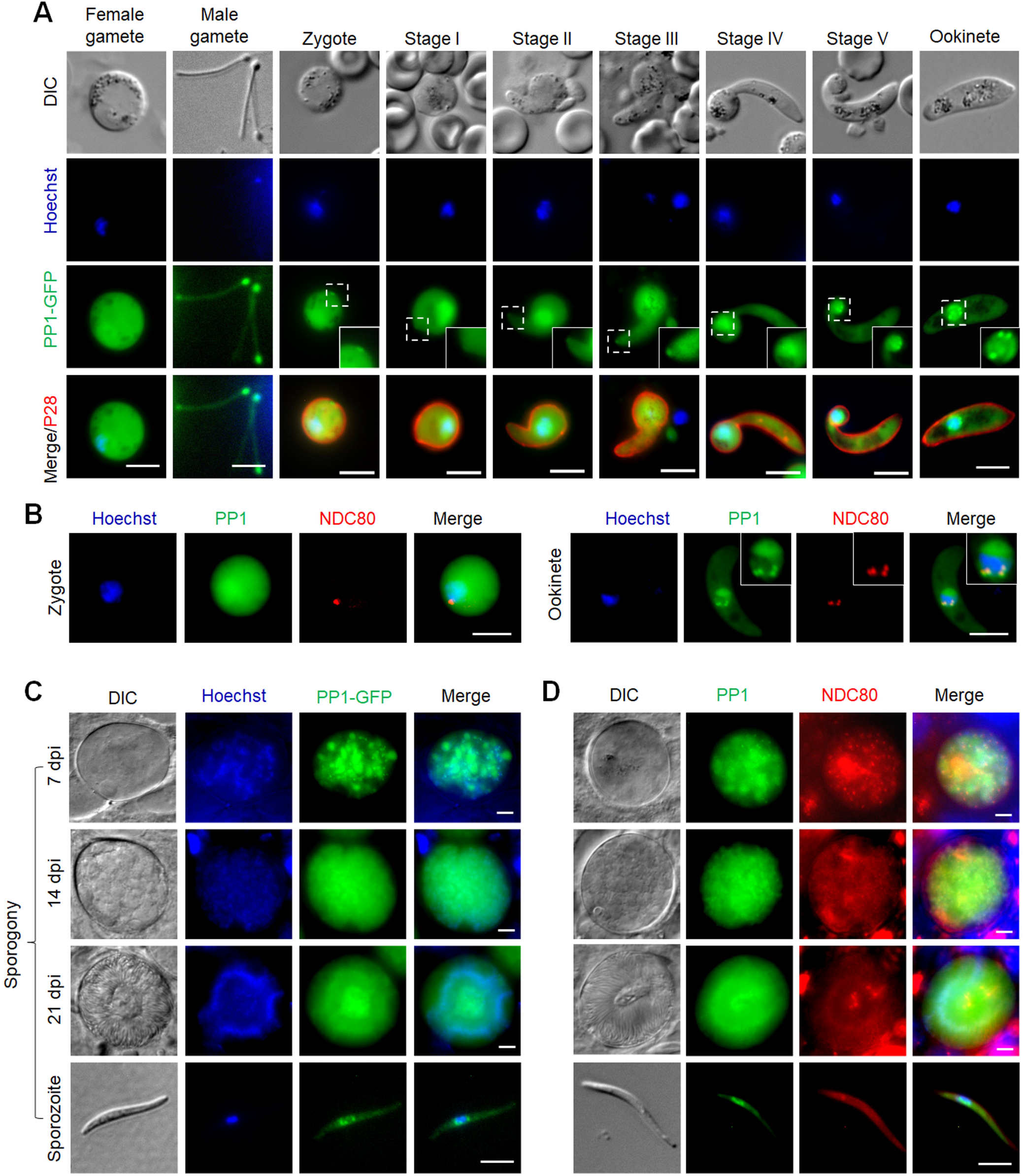
PP1-GFP localization during zygote formation, ookinete development, and sporogony inside the mosquito gut. **(A)** Live cell imaging showing PP1-GFP location in male and female gametes, zygote and during ookinete development (stages I to V and mature ookinete). A cy3-conjugated antibody, 13.1, which recognises the P28 protein on the surface of zygotes and ookinetes, was used to mark these stages. Panels: DIC (differential interference contrast), PP1-GFP (green, GFP), 13.1 (red), Merged: Hoechst (blue, DNA), PP1-GFP (green, GFP) and P28 (red). Scale bar = 5 μm higher magnification, the PP1-GFP signal on the zygote and developing apical end of early stage ‘retorts’, and in the nucleus of late retorts and ookinete stage. **(B)** Live cell imaging of PP1-GFP (green) in relation to NDC80-mCherry (red) and Hoechst staining (blue, DNA) in zygote and ookinete stages. **(C)** Live cell imaging of PP1-GFP in developing oocysts in mosquito guts at 7-, 14- and 21-days post-infection and in a sporozoite. Panels: DIC, Hoechst (blue, DNA), PP1-GFP (green), Merged (blue and green channels). **(D)** Live cell imaging of PP1-GFP in relation to NDC80 in developing oocysts and in a sporozoite. Panels: DIC (differential interference contrast), PP1-GFP (green), NDC80-mCherry (red), Merge (Hoechst, blue, DNA; PP1-GFP, green; NDC80-mCherry, red). Scale bar = 5 µm

### PP1-GFP has a diffuse distribution with multiple nuclear foci during oocyst development, representing endomitosis

Upon maturation the ookinete penetrates the mosquito midgut wall and embeds in the basal lamina to form an oocyst. Over the course of 21 days sporogony occurs in which several rounds of closed endomitotic division produce a multiploid cell termed a sporoblast^44^; with subsequent nuclear division resulting in thousands of haploid nuclei and concomitant sporozoite formation. We observed multiple foci of PP1GFP along with diffused localization during oocyst development (**Fig 3C**). Dual colour imaging showed a partial co-localisation of nuclear PP1-GFP and NDC80-mCherry foci throughout oocyst development and in sporozoites (**Fig 3D**), suggesting that PP1-GFP is diffusely distributed during oocyst development but also recruited to kinetochores, as highlighted by multiple nuclear foci during oocyst development.

### Generation of transgenic parasites with conditional knockdown of PP1

Our previous systematic analysis of the *P. berghei* protein phosphatases suggested that PP1 is likely essential for erythrocytic development^23^, a result which was further substantiated in a recent study of *P. falciparum*^30^. Since little is known about the role of PP1 in cell division during the sexual stages of parasite development in the mosquito vector, we attempted to generate transgenic, conditional knockdown lines using either the auxin inducible degron (AID) or the promoter trap double homologous recombination (PTD) systems. Despite successful generation of a parasite line expressing PP1-AID, addition of indole-3-acetic acid (IAA) did not result in PP1 depletion (**Fig S2C**) and male gametogony was not affected (**Fig S2C**). Therefore, we used a promoter trap method that had been used previously for functional analysis of the condensin core subunits SMC2/4 and the APC/C complex component, APC3^45, 46^. Double homologous recombination was used to insert the *ama* promoter upstream of *pp1* in a parasite line that constitutively expresses GFP^47^. The expected outcome was a parasite expressing PP1 during asexual blood stage development, but not at high levels during the sexual stages (**Fig S2D**). This strategy resulted in the successful generation of two independent PP1PTD parasite clones (clones 3 and 5) produced in independent transfections, and integration confirmed by diagnostic PCR (**Fig S2E**). The PP1PTD clones had the same phenotype and data presented here are combined results from both clones. Quantitative real time PCR (qRT-PCR) confirmed a downregulation of *pp1* mRNA in gametocytes by approximately 90% (**Fig 4A**).

**Fig. 4.**
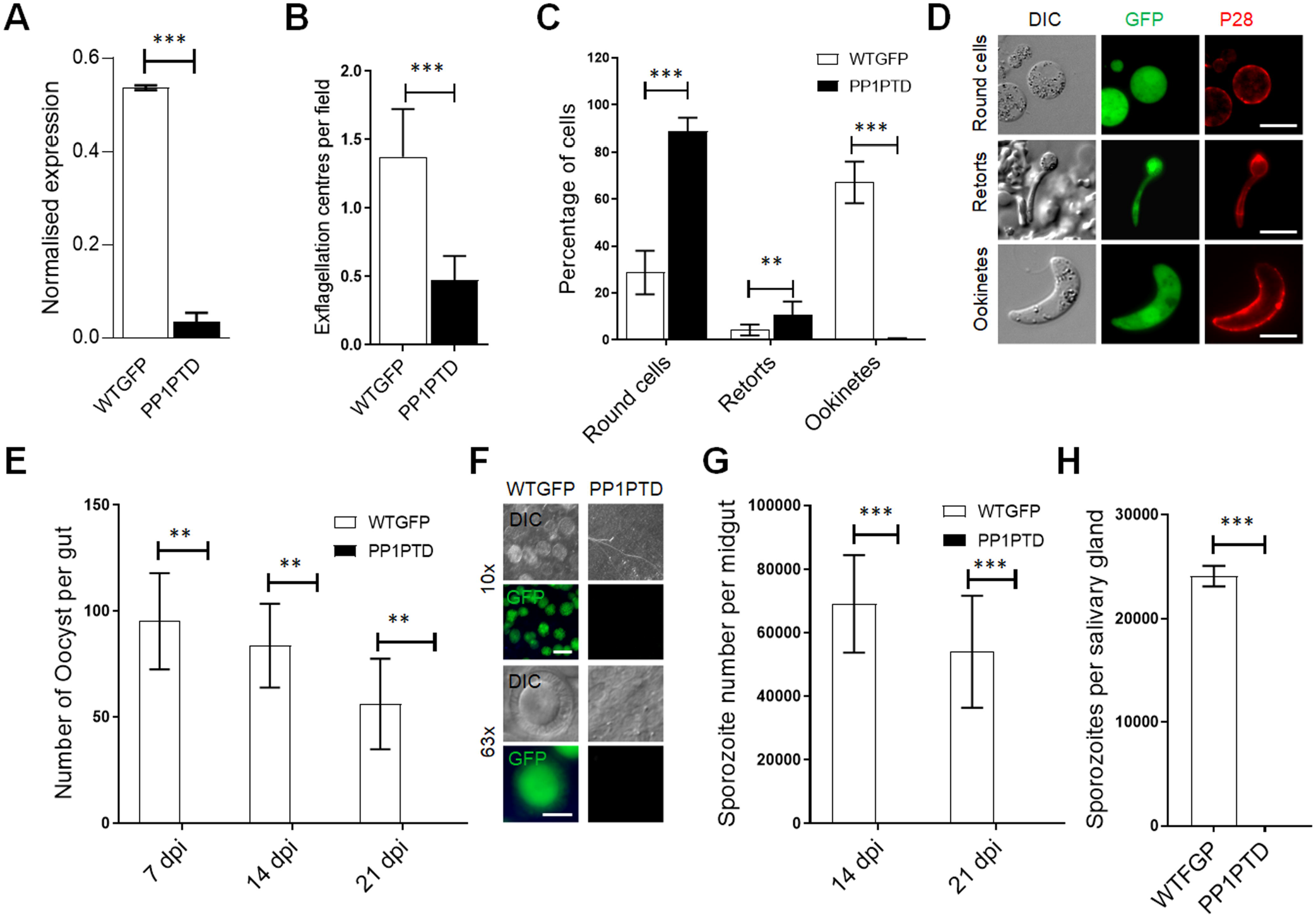
PP1 has an important role during male gamete formation and zygote - ookinete development. **(A)** qRT-PCR analysis of *pp1* transcription in PP1PTD and WT-GFP parasites, showing the downregulation of *pp1*. Each bar is the mean of three biological replicates ± SD. **(B)** Male gametogony (exflagellation) of PP1PTD line (black bar) and WT-GFP line (white bar) measured as the number of exflagellation centres per field. Mean ± SD; n=3 independent experiments. **(C)** Ookinete conversion as a percentage for PP1PTD and WT-GFP parasites. Ookinetes were identified using P28 antibody as a surface marker and defined as those cells that differentiated successfully into elongated ‘banana shaped’ ookinetes. Round cells show zygotes that did not start to transform and ‘retorts’ could not differentiate successfully in ookinetes. Mean ± SD; n=3 independent experiments. **(D)** Representative images of round cells, retorts and fully differentiated ookinetes. **(E)** Total number of GFP-positive oocysts per infected mosquito in PP1PTD and WT-GFP parasites at 7-, 14- and 21-days post infection (dpi). Mean ± SD; n=3 independent experiments. **(F)** Representative images of mosquito midguts on day 14 showing them full of oocysts in WTGFP and no oocyst in PP1PTD **(G)** Total number of sporozoites in oocysts of PP1PTD and WT-GFP parasites at 14 and 21 dpi. Mean ± SD; n=3 independent experiments. **(H)** Total number of sporozoites in salivary glands of PP1PTD and WT-GFP parasites. Bar diagram shows mean ± SD; n=3 independent experiments. Unpaired t-test was performed for statistical analysis. *p<0.05, **p<0.01, ***p<0.001.

### Knockdown of PP1 gene expression during the sexual stages blocks parasite transmission by affecting parasite growth and development within the mosquito vector

The PP1PTD parasites grew slower than the WT-GFP controls in the asexual blood stage (**Fig S2E**) but produced normal numbers of gametocytes in mice injected with phenylhydrazine before infection. Gametocyte activation with xanthurenic acid in ookinete medium resulted in significantly fewer exflagellating PP1PTD parasites per field in comparison to WT-GFP parasites, suggesting that male gamete formation was severely retarded (**Fig. 4B**). For those few PP1PTD gametes that were released and viable, zygote-ookinete differentiation following fertilization was severely affected (**Fig 4C**), with significantly reduced numbers of fully formed, banana-shaped ookinetes. In all PP1PTD samples analysed 24 hours post-fertilisation, the vast majority of zygotes were still round, and there was a significant number of abnormal retort-shaped cells with a long thin protrusion attached to the main cell body (**Fig 4C, D**).

To investigate further the function of PP1 during parasite development in the mosquito, *Anopheles stephensi* mosquitoes were fed on mice infected with PP1PTD and WT-GFP parasites as a control. The number of GFP-positive oocysts on the mosquito gut wall was counted on days 7, 14 and 21 post-infection. No oocysts were detected from PP1PTD parasites; whereas WT-GFP lines produced normal oocysts (**Fig 4E, F**), all of which had undergone sporogony (**Fig 4G**). Furthermore, no sporozoites were observed in the salivary glands of PP1PTD parasite-infected mosquitoes, in contrast to WTGFP parasite-infected mosquitoes (**Fig 4H**). This lack of viable sporozoites was confirmed by bite back experiments that showed no further transmission of PP1PTD parasites, in contrast to the WTGFP lines, which showed positive blood stage infection four days after mosquito feed (**Fig S2F**).

### Ultrastructure analysis of PP1PTD gametocytes shows defects in nuclear pole and axoneme assembly

To determine the ultrastructural consequences of reduced PP1 expression, WTGFP and PP1PTD gametocytes were examined at 6- and 30-min post activation (pa) by electron microscopy. At 6 min pa, the WTGFP male gametocytes exhibited an open nucleus with a number of nuclear poles (**Fig 5a**). Basal bodies and normal 9+2 axonemes were often associated with the nuclear poles (**Fig 5b,c**). At 6 min pa, the PP1PTD line exhibited a similar morphology (**Fig 5d-f**). However, quantitative analysis showed that PP1PTD gametocytes were relatively less well-developed, with fewer nuclear poles (0.86 WTGFP v. 0.56 PP1PTD), basal bodies (1.03 WTGFP v. 0.56 PP1PTD) and axonemes (2.57 WTGFP v. 1.27 PP1PTD), based on random sections of fifty male gametocytes in each group.

**Figure 5:**
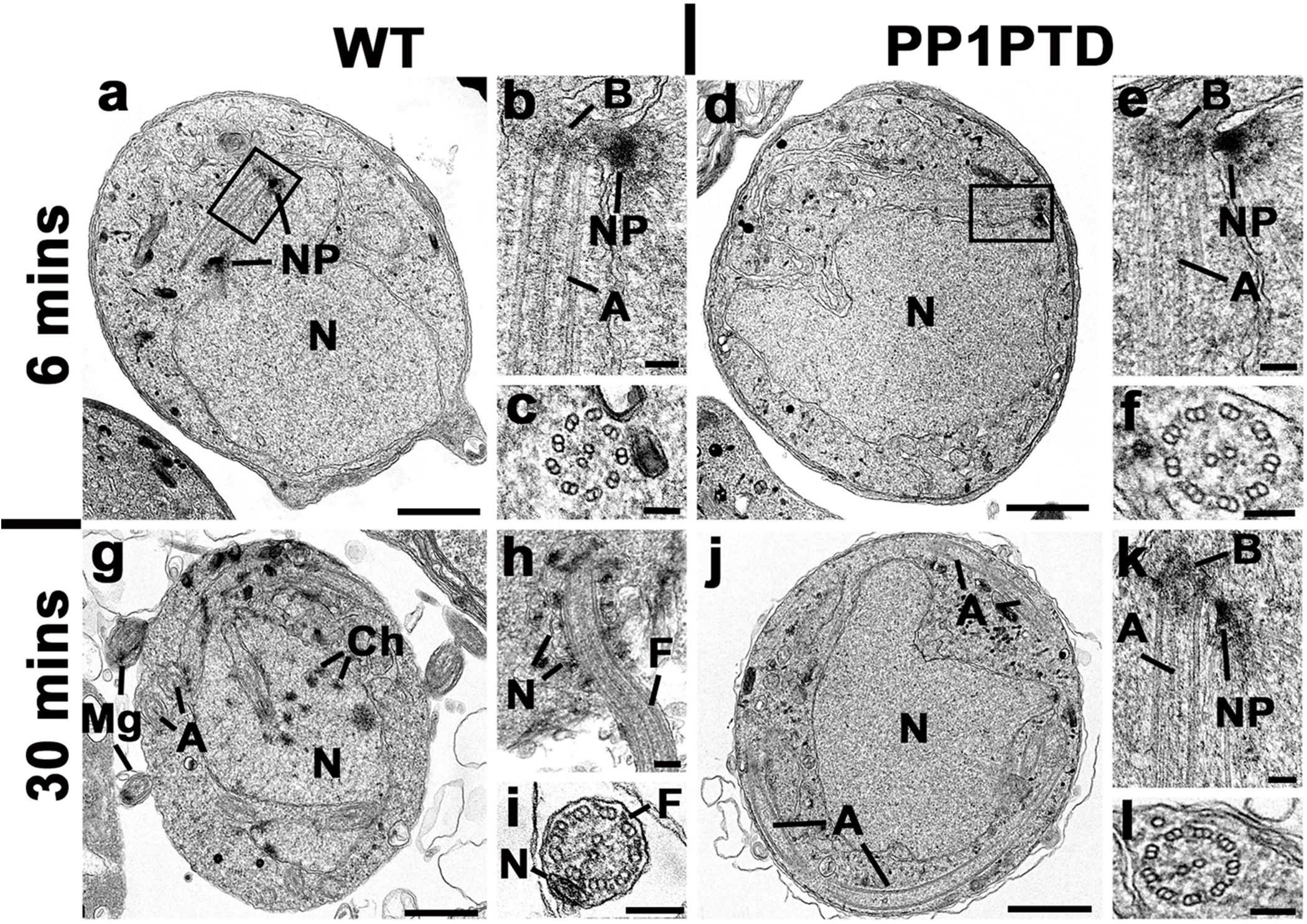
Ultrastructure analysis of PP1PTD gametocytes shows defects in nuclear pole and axoneme assembly during male gametogony. Electron micrographs of WTGFP (a-c, g-i) and PP1PTD (d-f, j-l) male gametocytes at 6 min (a-f) and 30 min (g-i) post activation (pa). Bars represent 1 µm (a, d, g, j) and 100 nm in all other micrographs. **(a)** Low power micrograph of a WTGFP male gametocyte with two nuclear poles (NP) associated with the nucleus (N). **(b)** Enlargement of the enclosed area showing the nuclear pole (NP) with adjacent basal body (B) and associated axoneme (A). **(c)** Cross section of an axoneme showing the 9+2 microtubule arrangement. **(d)** Low power micrograph of PP1PTD male gametocyte showing the central nucleus (N) with a nuclear pole and associated basal body (enclosed area). **(e)** Enlargement of the enclosed area showing the nuclear pole (NP) with adjacent basal body (B) and associated axoneme (A). **(f)** Cross section through an axoneme showing the 9+2 arrangement of microtubules. **(g)** Low power micrograph of a 30 min pa WTGFP male gametocyte showing the nucleus with areas of condensed chromatin. Note the cross sectioned free male gametes (Mg). A – axoneme. **(h)** Periphery of a male gametocyte undergoing exflagellation with the nucleus (N) associated with flagellum (F) protruding from the surface. **(i)** Cross section of a free male gamete showing the 9+2 microtubules of the flagellum (F) and electron dense nucleus (N). **(j)** Low power micrograph of a PP1PTD male gametocyte at 30 min pa showing the central nucleus (N) with a nuclear pole and an increased number of axoneme profiles (A) within the cytoplasm. **(k)** Detail of the periphery of a nucleus showing the nuclear pole (NP), basal body (B) and associated axoneme (A). **(l)** Cross section of an axoneme showing the 9+2 microtubule arrangement.

At 30 min post activation, most WTGFP male gametocytes (83%) were at a late stage of development with nuclei showing chromatin condensation (**Fig 5g**) and examples of exflagellation with the flagellate gamete protruding from the surface (**Fig 5h**) and free male gametes (**Fig 5g, i**). In contrast, most PP1PTD male gametocytes (85%) were stalled at an early stage of development similar to that at 6 min pa (**Fig 5j-l, Fig S3A**), based on random sections of fifty male gametocytes in each group. The main morphological difference between the two parasite lines was a marked increase in the number and length of the axonemes in the PP1PTD parasites at 30 min pa **(cf** **Figs 5a** **and j)**. In summary, the development of PP1PTD male gametocytes was severely retarded, although axoneme growth increased.

### PP1PTD parasites have altered expression of genes involved in cell cycle progression, cell motility, and apical cell polarity

To determine the consequences of PP1 knockdown on global mRNA transcription, we performed RNA-seq in duplicate on PP1PTD and WT-GFP gametocytes, 0 min and 30 min post-activation (**Fig 6A** **and B**). All replicates were clustered together based on condition (**Fig S3B**) and totals of 13 to 32 million RNA-seq reads were generated per sample for the global transcriptome **(Fig S3C).** In both 0 min and 30 min activated gametocytes, *pp1* was significantly down-regulated (by more than 16- fold, q value < 0.05) in PP1PTD parasites compared to WT-GFP parasites (**Fig 6A**), thus confirming the qPCR result (**Fig 4A**). We observed significant transcriptional perturbation in both 0 min and 30 min activated gametocytes, affecting the expression of 530 and 829 genes respectively (representing ∼ 10% and 16% of the total gene complement) (**Table S1**). Of the total perturbed genes, 344 and 581 were significantly down-regulated and 186 and 248 genes were significantly up-regulated in the PP1PTD parasites compared to WT-GFP controls in 0 min and 30 min activated gametocytes, respectively (**Table S1**).

**Figure 6:**
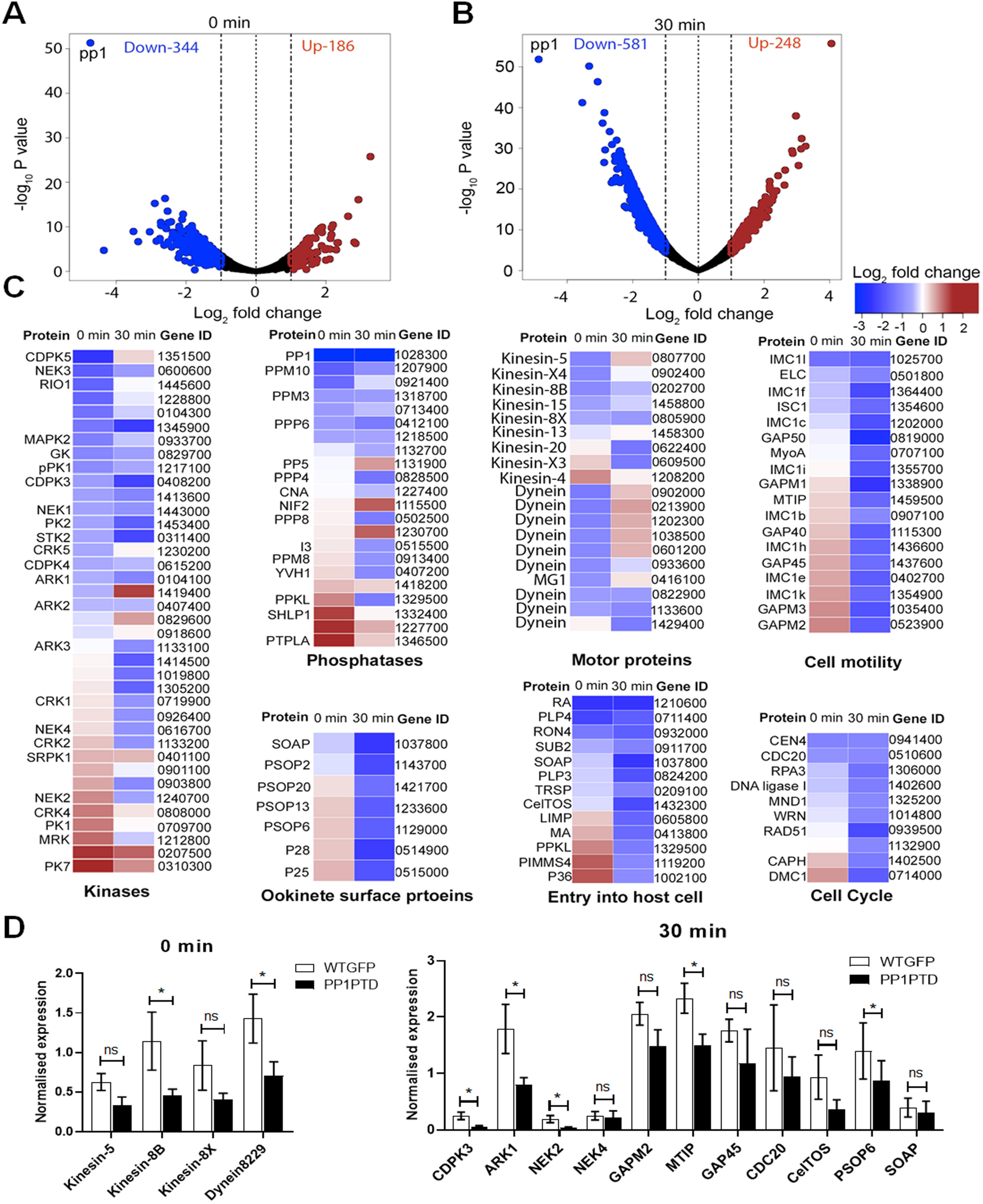
Transcriptome of PP1PTD mutant reveals important roles of PP1 in parasite cell cycle, motor protein function, and cell polarity during gametocyte biology. Volcano plots showing significantly down-regulated (blue Log_2_ fold ≤ -1, q value < 0.05) and up-regulated genes (brown Log_2_ fold ≥ 1, q value < 0.05) in PP1PTD compared to wild-type lines in non (0 min)-activated gametocytes **(A)** and 30 min activated gametocytes **(B).** Non-differentially regulated genes are represented as black dots. **(C)** Expression heat maps showing affected genes from specific functional classes of proteins such as kinases, phosphatases, motor proteins and proteins associated with parasite motility, entry into the host cell, ookinete surface and cell cycle. Genes are ordered based on their differential expression pattern in non-activated gametocytes. **(D)** Validation by qRT-PCR of a few genes randomly selected based on the RNA-seq data. All the experiments were performed three times in duplicate with two biological replicates. *p ≤05.

To explore the biological roles of the genes down-regulated following reduced *pp1* expression, we performed gene-ontology (GO) based enrichment analysis. We observed that many genes encoding kinases, phosphatases and motor proteins were differentially expressed in either or in both 0 min or 30 mins activated gametocytes (**Fig 6C**), complementing the observations from our phenotypic analysis (**Fig 4C, D**). In addition, we also observed that many genes encoding proteins involved in cell motility, cell-cycle progression, host-cell entry and ookinete development were significantly affected in activated PP1PTD gametocytes.

Transcript levels measured by RNA-seq were further validated by qRT-PCR for a few selected genes modulated in gametocytes at 0 min and 30 min, and involved in regulation of cell cycle, cell motility, and ookinete invasion (**Fig 6D**).

### PP1–GFP interacts with similar proteins in schizonts and gametocytes, but with a predominance of microtubule motor kinesins in gametocytes

Previous studies have analysed the PP1 interactome in *P. falciparum* schizonts, revealing several interacting partners^32, 48^. Here, we analysed the PP1 interactome in *P. berghei* schizonts and gametocytes to establish whether there were differences that might reflect distinct functions in the two stages. We immunoprecipitated PP1-GFP from lysates of schizonts following parasite culture *in vitro* for 10 hours and 24 hours and from lysates of gametocytes 10 to 11 min post activation because of high PP1-GFP abundance at these stages (**Fig 7A**, **Table S2**). Mass spectrometric analysis of these pulldowns identified several proteins common to both schizont and gametocyte stages, suggesting similar protein complexes in both stages (**Fig 7A, B**). In addition, we also identified in the pulldown from gametocyte lysates a number of microtubule proteins that are associated with the spindle or axoneme, including kinesin-8B, kinesin-15, kinesin-13 and PF16 (**Fig 7A, B**). These proteins are specific to male gametocytes and may have important roles in axoneme assembly and male gametogony^43, 49^.

**Fig. 7.**
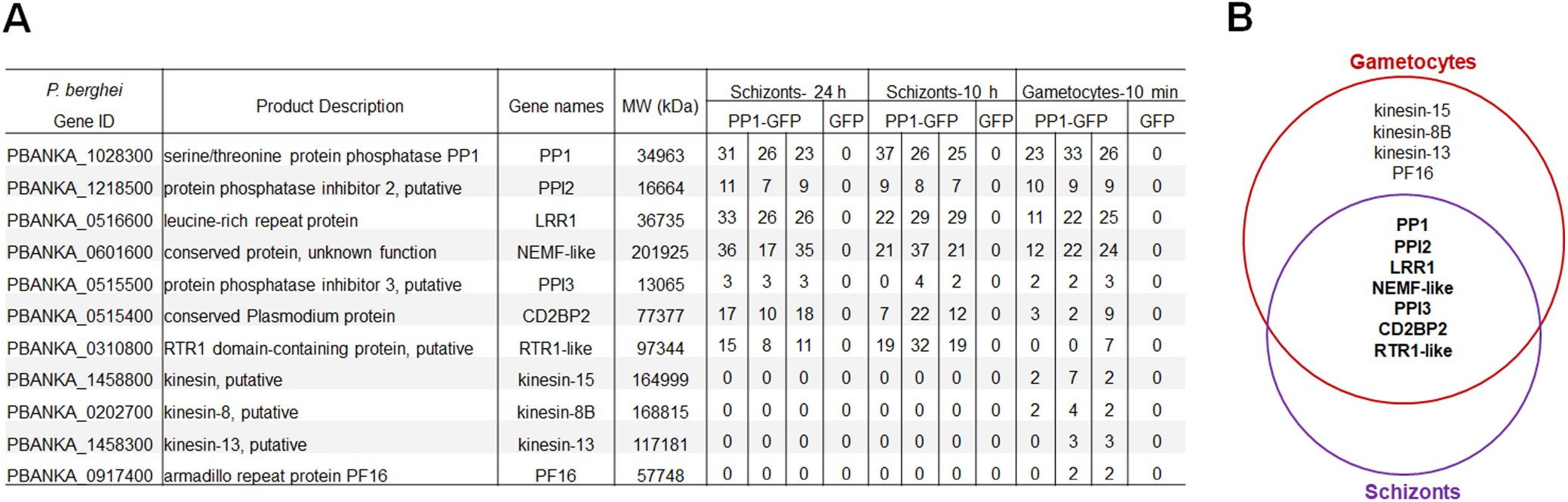
Interacting partners of PP1 during asexual schizont and sexual gametocyte stages. **(A)** List of proteins interacting with PP1 during schizont and gametocyte stages. **(B)** Venn diagram showing common interacting partners in schizonts and gametocytes with some additional proteins specific to gametocytes.

## Discussion

Reversible protein phosphorylation is crucial for cell cycle progression in eukaryotes and is tightly controlled by a variety of protein kinases and phosphatases^50, 51^. *Plasmodium* possesses a set of divergent protein kinases and protein phosphatases, which regulate many processes during cell division and parasite development throughout different stages of the life-cycle^23, 52^, and PP1 is quantitatively one of the most important protein phosphatases that hydrolyse serine/threonine linked phosphate ester bonds^53^. It is expressed in all cells and is highly conserved in organisms throughout the Eukaryota including *Plasmodium*^23, 54^, although a key region at the C-terminus that is known to regulate mitotic exit is missing from the parasite PP1^24^. It forms stable complexes with several PP1-interacting proteins (PIPs) that are diverse and differ in various organisms, control its location and function, and assist in processes throughout the cell cycle^8, 55, 56, 57^.

The cell cycle in *Plasmodium* differs from that of many other eukaryotes with atypical mitotic and meiotic divisions throughout its life cycle^41, 43, 58^. It lacks several classical cell cycle regulators including the protein phosphatases CDC14 and CDC25^23^, and key protein kinases including polo-like kinases^52, 59^. Therefore, the role of PP1, or indeed other PPPs in the absence of these protein phosphatases and kinases may be more crucial during cell division in *Plasmodium*. In the present study, we focused on the location and functional role of PP1 in mitotic division, particularly during male gametogony, and in meiosis during the zygote to ookinete transition. Both processes are essential for the sexual stage of the *Plasmodium* life cycle and are required for transmission by the mosquito vector. Our previous systematic functional analysis of the *Plasmodium* protein phosphatome^23^ showed that PP1 has an essential role during asexual blood stage development; this finding was supported by a recent study, which showed its role in merozoite egress from the host erythrocyte^30^. Our findings here reveal that PP1 is also expressed constitutively and co-localises with the kinetochore protein NDC80 during different stages of *Plasmodium* asexual and sexual development, hinting at a role during atypical chromosome segregation. Our conditional PP1 gene knockdown suggests that it plays a crucial role in mitotic division during male gametogony; and may regulate cell polarity in meiosis during zygote to ookinete transformation.

Male gametogony in *Plasmodium* is a rapid process with DNA replication and three mitotic divisions followed by karyokinesis and cytokinesis to form eight flagellated male gametes within 15 min of gametocyte activation^41, 58^. The expression and localization profiles show that although the protein is diffusely distributed throughout the cytoplasm and the nucleus, there is a cyclic enrichment of PP1-GFP at the kinetochore, associating with NDC80-mCherry during all three successive genome replication and closed mitotic divisions without nuclear division^41^. The accumulation of PP1-GFP at the kinetochore at the start of nuclear division and subsequent decrease upon completion is similar to the situation in other eukaryotes where the activity of PP1 increases in G2 phase, reduces during prophase and metaphase, and increases again during anaphase^51^. This behaviour suggests a role for PP1 in regulating rapid mitotic entry and exit during male gametogony. Further analysis of PP1 gene knockdown showed a significant decrease in male gamete formation (i.e. during male gametogony), and ultrastructural analysis revealed fewer nuclear poles and basal bodies associated with axonemes, with a concomitant absence of chromosome condensation in male gametocytes of PP1PTD parasites, suggesting a role for PP1 during chromosome segregation and gamete formation (i.e. flagella formation). A similar phenotype was observed in our recent study of a divergent *Plasmodium* cdc2-related kinase (CRK5)^60^. CRK5 deletion resulted in fewer nuclear poles, and no chromatin condensation, cytokinesis or flagellum formation, suggesting that there may be a coordinated activity of PP1 and CRK5 in reversible phosphorylation. A recent phosphoregulation study across the short time period of male gametogony showed tightly controlled phosphorylation events mediated by several protein kinases including ARK2, CRK5, and NEK1^61^; however, the reciprocal protein phosphatases that reverse these events are unknown. The expression pattern and cyclic enrichment of PP1 at the kinetochores suggest a key reciprocal role of PP1 in reversing the protein phosphorylation mediated by these kinases. This interpretation is supported by our global transcriptome analysis of the PP1PTD parasite, which showed a modulated expression of several serine/threonine protein kinases, protein phosphatases and motor proteins. The similar pattern of reduced expression of some of these protein phosphatases and kinases suggests they have a coordinated role in reversible phosphorylation. Of note, phosphoregulation of motor proteins in *Plasmodium* has been described previously^61^ and it plays an important role in spindle assembly and axoneme formation during male gametogony^41, 43^.

The proteomics analysis showed that PP1 interacts with a conserved set of proteins in both asexual and sexual stages of development, and with additional proteins in gametocytes, implicating major motor proteins that may be required for spindle and axoneme assembly during male gametogenesis. Several of the conserved interacting partners also form complexes with PP1 in other eukaryotes but the gametocyte specific proteins such as kinesin-8B, kinesin-15, kinesin-13, and PF16 are unique PIPs in *Plasmodium*^9, 48^. These findings are consistent with our transcriptomic analysis of PP1PTD showing modulated expression of motor proteins in male gametocytes.

Analysis of PP1 during the zygote to ookinete transformation showed an additional location at the nascent apical end during the early stages of ookinete differentiation, which may help define the cell’s polarity. This suggestion is supported by the consequence of PP1PTD gene knock down in which numerous underdeveloped ookinetes with a long, thin protrusion attached to the main cell body were observed. This idea is also substantiated by a recent study showing that apical-basal polarity in *Drosophila* is controlled by PP1-mediated, and SDS22-dependent, dephosphorylation of LGL, an actomyosin-associated protein ^62^. However, it is important to note that the suggested role in cell polarity is based on the observation that after fertilization only a few abnormal ookinetes are formed, which have the morphologically distinct elongated apical end. Our transcriptomic analysis of PP1PTD-gametocytes showed modulation of genes for several organelle markers such as CTRP and SOAP^63, 64^, polarity markers such as SAS6L and IMC proteins^65, 66^, as well as genes involved in gliding motility including other IMC proteins, MyoA, GAPs, GAPMs, and some ookinete specific proteins. These proteins are important for maintenance of cell shape, gliding motility, and mosquito gut wall invasion by ookinetes. These results suggest that PP1 may be involved in the phospho-regulation of proteins involved in defining polarity and maintenance of ookinete shape.

In conclusion, PP1 is a constitutively expressed phosphatase, distributed throughout the cell but enriched in the nucleus, and associated with the kinetochore during mitosis and meiosis, with a role in the regulation of mitosis and meiosis throughout the *Plasmodium* life-cycle.

## Material and Methods

### Ethics statement

The animal work performed in the UK passed an ethical review process and was approved by the United Kingdom Home Office. Work was carried out under UK Home Office Project Licenses (30/3248 and PDD2D5182) in accordance with the United Kingdom ‘Animals (Scientific Procedures) Act 1986’. Six- to eight-week-old female CD1 outbred mice from Charles River laboratories were used for all experiments in the UK.

### Generation of transgenic parasites

GFP-tagging vectors were designed using the p277 plasmid vector and transfected as described previously^23^. A schematic representation of the endogenous *pp1* locus (PBANKA_1028300), the constructs and the recombined *pp1* locus can be found in **Fig S1A**. For GFP-tagging of PP1 by single crossover homologous recombination, a region of *pp1* downstream of the ATG start codon was used to generate the construct. For the genotypic analyses, a diagnostic PCR reaction was performed as outlined in **Fig. S1A**. Primer 1 (intP6tg) and primer 2 (ol492) were used to determine correct integration of the *gfp* sequence at the targeted locus. For western blotting, purified gametocytes were lysed using lysis buffer (10 mM TrisHCl pH 7.5, 150 mM NaCl, 0.5 mM EDTA and 1% NP-40). The lysed samples were boiled for 10 min at 95 °C after adding Laemmli buffer and were centrifuged at maximum speed (13000g) for 5 min. The samples were electrophoresed on a 4–12% SDS-polyacrylamide gel. Subsequently, resolved proteins were transferred to nitrocellulose membrane (Amersham Biosciences). Immunoblotting was performed using the Western Breeze Chemiluminescence Anti-Rabbit kit (Invitrogen) and anti-GFP polyclonal antibody (Invitrogen) at a dilution of 1:1250, according to the manufacturer’s instructions.

To study the function of PP1, we used two conditional knock down systems; a <u>p</u>romoter exchange/trap using <u>d</u>ouble homologous recombination (PP1PTD) and an auxin inducible degron (PP1AID) system. The PP1AID construct was derived from the p277 plasmid, where the GFP sequence was excised following digestion with *Age*I and *Not*I restriction enzymes and replaced with an AID/HA coding sequence. The AID-HA sequence was PCR amplified (using primers: 5’-CCCCAGACGTCGGATCCAATGATGGGCAGTGTCGAGCT-3’ and 5’-ATATAAGTAAGAAAAACGGCTTAAGCGTAATCTGGA-3’) from the GW-AID/HA plasmid (http://plasmogem.sanger.ac.uk/). Fragments were assembled following the Gibson assembly protocol to generate the PP1-AID/HA transfection plasmid that was transfected in the 615 line. Conditional degradation of PP1-AID/HA was performed as described previously^67^. A schematic representation of the endogenous *pp1* locus (PBANKA_1028300), the constructs and the recombined *pp1* locus can be found in **Fig S2A.** A diagnostic PCR was performed for *pp1* gene knockdown parasites as outlined in **Fig. S2A**. Primer 1 and Primer 3 were used to determine successful integration of the targeting construct at the 3’ gene locus **(Fig S2B)**. Primer 1 and Primer 2 were used as controls **(Fig S2B)**.

The conditional knockdown construct PP1-PTD was derived from *P_ama1_* (pSS368)) where *pp1* was placed under the control of the *ama1* promoter, as described previously^68^. A schematic representation of the endogenous *pp1* locus, the constructs and the recombined *pp1* locus can be found in **Fig S2D**. A diagnostic PCR was performed for *pp1* gene knockdown parasites as outlined in **Fig. S2D**. Primer 1 (5’-intPTD36) and Primer 2 (5’-intPTD) were used to determine successful integration of the targeting construct at the 5’ gene locus. Primer 3 (3’-intPTD) and Primer 4 (3’-intPTama1) were used to determine successful integration for the 3’ end of the gene locus **(Fig. S2E)**. All the primer sequences can be found in **Table S3**. *P. berghei* ANKA line 2.34 (for GFP-tagging) or ANKA line 507cl1 expressing GFP (for the knockdown construct) parasites were transfected by electroporation^47^.

### Purification of schizonts and gametocytes

Blood cells obtained from infected mice (day 4 post infection) were cultured for 11h and 24 h at 37°C (with rotation at 100 rpm) and schizonts were purified the following day on a 60% v/v NycoDenz (in PBS) gradient, (NycoDenz stock solution: 27.6% w/v NycoDenz in 5 mM Tris-HCl, pH 7.20, 3 mM KCl, 0.3 mM EDTA).

The purification of gametocytes was achieved using a protocol described previously^69^ with some modifications. Briefly, parasites were injected into phenylhydrazine treated mice and enriched by sulfadiazine treatment after 2 days of infection. The blood was collected on day 4 after infection and gametocyte-infected cells were purified on a 48% v/v NycoDenz (in PBS) gradient (NycoDenz stock solution: 27.6% w/v NycoDenz in 5 mM Tris-HCl, pH 7.20, 3 mM KCl, 0.3 mM EDTA). The gametocytes were harvested from the interface and activated.

### Live cell imaging

To examine PP1-GFP expression during erythrocyte stages, parasites growing in schizont culture medium were used for imaging at different stages (ring, trophozoite, schizont and merozoite) of development. Purified gametocytes were examined for GFP expression and localization at different time points (0, 1-15 min) after activation in ookinete medium^43^. Zygote and ookinete stages were analyzed throughout 24 h of culture. Images were captured using a 63x oil immersion objective on a Zeiss Axio Imager M2 microscope fitted with an AxioCam ICc1 digital camera (Carl Zeiss, Inc).

### Generation of dual tagged parasite lines

The PP1-GFP parasites were mixed with NDC80-cherry and MyoA-cherry parasites in equal numbers and injected into mice. Mosquitoes were fed on mice 4 to 5 days after infection when gametocyte parasitaemia was high. These mosquitoes were checked for oocyst development and sporozoite formation at day 14 and day 21 after feeding. Infected mosquitoes were then allowed to feed on naïve mice and after 4 - 5 days, and the mice were examined for blood stage parasitaemia by microscopy with Giemsa-stained blood smears. In this way, some parasites expressed both PP1-GFP and NDC80-cherry; and PP1-GFP and MyoA-cherry in the resultant gametocytes, and these were purified and fluorescence microscopy images were collected as described above.

### Parasite phenotype analyses

Blood containing approximately 50,000 parasites of the PP1PTD line was injected intraperitoneally (i.p.) into mice to initiate infections. Asexual stages and gametocyte production were monitored by microscopy on Giemsa-stained thin smears. Four to five days post infection, exflagellation and ookinete conversion were examined as described previously ^70^ with a Zeiss AxioImager M2 microscope (Carl Zeiss, Inc) fitted with an AxioCam ICc1 digital camera. To analyse mosquito transmission, 30– 50 *Anopheles stephensi* SD 500 mosquitoes were allowed to feed for 20 min on anaesthetized, infected mice with an asexual parasitaemia of 15% and a comparable number of gametocytes as determined on Giemsa-stained blood films. To assess mid-gut infection, approximately 15 guts were dissected from mosquitoes on day 14 post feeding, and oocysts were counted on an AxioCam ICc1 digital camera fitted to a Zeiss AxioImager M2 microscope using a 63x oil immersion objective. On day 21 post-feeding, another 20 mosquitoes were dissected, and their guts crushed in a loosely fitting homogenizer to release sporozoites, which were then quantified using a haemocytometer or used for imaging. Mosquito bite back experiments were performed 21 days post-feeding using naive mice, and blood smears were examined after 3-4 days.

### Electron microscopy

Gametocytes activated for 6 min and 30 min were fixed in 4% glutaraldehyde in 0.1 M phosphate buffer and processed for electron microscopy as previously described^71^. Briefly, samples were post fixed in osmium tetroxide, treated *en bloc* with uranyl acetate, dehydrated and embedded in Spurr’s epoxy resin. Thin sections were stained with uranyl acetate and lead citrate prior to examination in a JEOL JEM-1400 electron microscope (JEOL Ltd, UK)

### Quantitative Real Time PCR (qRT-PCR) analyses

RNA was isolated from gametocytes using an RNA purification kit (Stratagene). cDNA was synthesised using an RNA-to-cDNA kit (Applied Biosystems). Gene expression was quantified from 80 ng of total RNA using a SYBR green fast master mix kit (Applied Biosystems). All the primers were designed using the primer3 software (https://primer3.ut.ee/). Analysis was conducted using an Applied Biosystems 7500 fast machine with the following cycling conditions: 95°C for 20 s followed by 40 cycles of 95°C for 3 s; 60°C for 30 s. Three technical replicates and three biological replicates were performed for each assayed gene. The *hsp70* (PBANKA_081890) and *arginyl-t RNA synthetase* (PBANKA_143420) genes were used as endogenous control reference genes. The primers used for qPCR can be found in **Table S1**.

### Transcriptome study using RNA-seq

For RNA extraction, parasite samples were passed through a plasmodipur column to remove host DNA contamination prior to RNA isolation. Total RNA was extracted from activated gametocytes and schizonts of WT-GFP and PP1PTD parasites (two biological replicates each) using an RNeasy purification kit (Qiagen). RNA was vacuum concentrated (SpeedVac) and transported using RNA-stable tubes (Biomatrica). Strand-specific 354mRNA sequencing was performed on total RNA and using TruSeq stranded mRNA sample prep 355kit LT (Illumina), as previously described^72^. Libraries were sequenced using an Illumina Hiseq 4000 sequencing platform with paired-end 150 bp read chemistry. The quality of the raw reads was assessed using FATSQC (http://www.bioinformatics.babraham.ac.uk/projects/fastqc). Low-quality reads and Illumina adaptor sequences from the read ends were removed using Trimmomatic R^73^. Processed reads were mapped to the *P. beghei ANKA* reference genome (release 40 in PlasmoDB - http://www.plasmoddb.org) using Hisat2^74^ (V 2.1.0) with parameter “—rna-strandness FR”. Counts per feature were estimated using FeatureCounts^75^. Raw read counts data were converted to counts per million (cpm) and genes were excluded if they failed to achieve a cpm value of 1 in at least one of the three replicates performed. Library sizes were scale-normalized by the TMM method using EdgeR software^76^ and further subjected to linear model analysis using the voom function in the limma package^77^. Differential expression analysis was performed using DeSeq2^78^. Genes with a fold-change greater than two and a false discovery rate corrected p-value (Benjamini-Hochberg procedure) < 0.05 were considered to be differentially expressed. Functional groups shown in Figure 6C were inferred from annotations available in PlasmoDB: Release 49 (https://plasmodb.org/plasmo/app).

### Immunoprecipitation and Mass Spectrometry

Schizonts, following 11 hours and 24 hours, respectively in *in vitro* culture, and male gametocytes 11 min post activation were used to prepare cell lysates. Purified parasite pellets were crosslinked using formaldehyde (10 min incubation with 1% formaldehyde, followed by 5 min incubation in 0.125M glycine solution and 3 washes with phosphate buffered saline (PBS) (pH, 7.5). Immunoprecipitation was performed using crosslinked protein and a GFP-Trap^®^_A Kit (Chromotek) following the manufacturer’s instructions. Proteins bound to the GFP-Trap^®^_A beads were digested using trypsin and the peptides were analysed by LC-MS/MS. Briefly, to prepare samples for LC-MS/MS, wash buffer was removed, and ammonium bicarbonate (ABC) was added to beads at room temperature. We added 10 mM TCEP (Tris-(2-carboxyethyl) phosphine hydrochloride) and 40 mM 2-chloroacetamide (CAA) and incubation was performed for 5 min at 70°C. Samples were digested using 1 µg Trypsin per 100 µg protein at room temperature overnight followed by 1% TFA addition to bring the pH into the range of 3-4 before mass spectrometry.

### Statistical analysis

All statistical analyses were performed using GraphPad Prism 8 (GraphPad Software). An unpaired t-test and two-way anova test were used to examine significant differences between wild-type and mutant strains for qRT-PCR and phenotypic analysis accordingly.

## Data Availability

RNA Sequence reads have been deposited in the NCBI Sequence gene expression omnibus with the accession number GSE164175.

“The mass spectrometry proteomics data have been deposited to the ProteomeXchange Consortium with the dataset identifier PXD023571 and 10.6019/PXD023571.

## Author contributions

RT and MZ conceived and designed all experiments. RT, MZ, RP, DB, DSG, GK performed the GFP tagging and conditional knockdown with promoter trap experiments. RR and MBr generated and characterised the PP1-AID/HA line. MZ, RP, GK, DB and RT performed protein pull-down experiments. ARB performed mass spectrometry. AS, RN and AP performed RNA sequencing (RNA-seq). DJPF and SV performed electron microscopy. RT, AAH, MZ, DSG, AS, RN, AP and DJPF analyzed the data. MZ, DSG and RT wrote the original draft. RT, AAH, AP, DJPF, MB edited and reviewed the manuscript and all other contributed to it.

## Acknowledgments

We thank Julie Rodgers for helping to maintain the insectary and other technical works, and the personnel at the Bioscience Core Laboratory (BCL) in KAUST for sequencing the RNA samples and producing the raw datasets.

## Supplementary figures

**Fig. S1. Generation and genotypic analysis of PP1GFP parasites (A)** Schematic representation for 3’-tagging of *pp1* gene with green fluorescent protein (GFP) sequence via single homologous recombination. **(B)** Integration PCR showing correct integration of tagging construct. **(C)** Western blot showing expected size of PP1-GFP protein.

**Fig. S2. Generation and genotype analysis of conditional knockdown PP1 parasites**

**(A)** Schematic representation of auxin inducible degron (AID) strategy to generate PP1AID parasites. **(B)** Integration PCR of the PP1AID construct in the *pp1* locus. Primer 1 and Primer 2 were used for control PCR while primer 1 and primer 3 were used to determine successful integration of AID-HA sequence and selectable marker at 3’-end of *pp1* locus **(C)** PP1AID protein expression level as measured by western blotting upon addition of auxin to mature purified gametocytes; α-tubulin serves as a loading control. Auxin treatment of PP1AID showed no defect in exflagellation (error bars show standard deviation from the mean; technical replicates from three independent infections. **(D)** Schematic representation of the promoter swap strategy (PP1PTD, placing *pp1* under the control of the *ama1* promoter) by double homologous recombination. Arrows 1 and 2 indicate the primer positions used to confirm 5’ integration and arrows 3 and 4 indicate the primers used for 3’ integration. **(E)** Integration PCR of the promotor swap construct into the *pp1* locus. Primer 1 (5’-IntPTD36) with primer 2 (5’-IntPTD) were used to determine successful integration of the selectable marker. Primer 3 (3’-intPTama1) and primer 4 (3’-IntPTD36) were used to determine the successful integration of ama1 promoter. Primer 1 (5’-IntPTD36) and primer 4 (3’-IntPTD36) were used to show complete knock-in of the construct and the absence of a band at 2.1 kb (endogenous) resulting in complete knock-in of the construct. **(F)** Parasitaemia during blood stage schizogony showing a significant slow growth of PP1PTD compared to WTGFP parasites. Experiment was done with three mice each with 1000 parasites per mice injected intraperitonially. ***P<0.001 **(G)** Bite back experiments show no transmission of PP1PTD parasites (black bar) from mosquito to mouse, while successful transmission was shown by WT-GFP parasites. Mean ± SD; n= 3 independent experiments.

**Fig. S3. Analysis of PP1PTD development and RNA seq analysis: (A)** The quantification of electron microscopy data showing PP1PTD male gametocytes halted at an early stage of development in comparison with WTGFP male gametocytes at 30 min post activation. These data are based on analysis of fifty random sections of male gametocytes per sample.

**(B)** Clustered dendrogram of two biological replicates of WTGFP and PP1PTD mutant parasite lines during gametocyte stage using hierarchical clustering algorithm. Analysis was performed on normalized count data. **(C)** RNA-seq read statistics.

## Supplementary Tables

Table S1. Differentially expressed genes in PP1PTD parasites

Table S2. List of proteins pulled down by PP1GFP

Table S3. Primers used in this study

## References

1. Marston AL, Amon A. Meiosis: cell-cycle controls shuffle and deal. Nat Rev Mol Cell Biol 5, 983–997 (2004).

2. Sullivan M, Morgan DO. Finishing mitosis, one step at a time. Nat Rev Mol Cell Biol 8, 894–903 (2007).

3. Novak B, Kapuy O, Domingo-Sananes MR, Tyson JJ. Regulated protein kinases and phosphatases in cell cycle decisions. Curr Opin Cell Biol 22, 801–808 (2010).

4. Afshar K, Werner ME, Tse YC, Glotzer M, Gonczy P. Regulation of cortical contractility and spindle positioning by the protein phosphatase 6 PPH-6 in one-cell stage C. elegans embryos. Development 137, 237–247 (2010).

5. Boutros R, Dozier C, Ducommun B. The when and wheres of CDC25 phosphatases. Curr Opin Cell Biol 18, 185–191 (2006).

6. Chen F, et al. Multiple protein phosphatases are required for mitosis in Drosophila. Curr Biol 17, 293–303 (2007).

7. Stegmeier F, Amon A. Closing mitosis: the functions of the Cdc14 phosphatase and its regulation. Annu Rev Genet 38, 203–232 (2004).

8. Bollen M, Gerlich DW, Lesage B. Mitotic phosphatases: from entry guards to exit guides. Trends Cell Biol 19, 531–541 (2009).

9. Heroes E, Lesage B, Gornemann J, Beullens M, Van Meervelt L, Bollen M. The PP1 binding code: a molecular-lego strategy that governs specificity. FEBS J 280, 584–595 (2013).

10. Holder J, Poser E, Barr FA. Getting out of mitosis: spatial and temporal control of mitotic exit and cytokinesis by PP1 and PP2A. FEBS Lett 593, 2908–2924 (2019).

11. Nilsson J. Protein phosphatases in the regulation of mitosis. J Cell Biol 218, 395–409 (2019).

12. Hattersley N, et al. A Nucleoporin Docks Protein Phosphatase 1 to Direct Meiotic Chromosome Segregation and Nuclear Assembly. Dev Cell 38, 463–477 (2016).

13. Cheeseman IM. The kinetochore. Cold Spring Harb Perspect Biol 6, a015826 (2014).

14. Nagpal H, Fukagawa T. Kinetochore assembly and function through the cell cycle. Chromosoma 125, 645–659 (2016).

15. Varma D, Salmon ED. The KMN protein network--chief conductors of the kinetochore orchestra. J Cell Sci 125, 5927–5936 (2012).

16. Lampert F, Westermann S. A blueprint for kinetochores - new insights into the molecular mechanics of cell division. Nat Rev Mol Cell Biol 12, 407–412 (2011).

17. Wu JQ, et al. PP1-mediated dephosphorylation of phosphoproteins at mitotic exit is controlled by inhibitor-1 and PP1 phosphorylation. Nat Cell Biol 11, 644–651 (2009).

18. Dohadwala M, et al. Phosphorylation and inactivation of protein phosphatase 1 by cyclin-dependent kinases. Proc Natl Acad Sci U S A 91, 6408–6412 (1994).

19. Bouchoux C, Uhlmann F. A quantitative model for ordered Cdk substrate dephosphorylation during mitotic exit. Cell 147, 803–814 (2011).

20. Grallert A, et al. A PP1-PP2A phosphatase relay controls mitotic progression. Nature 517, 94–98 (2015).

21. Mochida S, Hunt T. Protein phosphatases and their regulation in the control of mitosis. EMBO Rep 13, 197–203 (2012).

22. Howick VM, et al. The Malaria Cell Atlas: Single parasite transcriptomes across the complete Plasmodium life cycle. Science 365, (2019).

23. Guttery David S, et al. Genome-wide Functional Analysis of Plasmodium Protein Phosphatases Reveals Key Regulators of Parasite Development and Differentiation. Cell Host & Microbe 16, 128–140 (2014).

24. Khalife J, Freville A, Gnangnon B, Pierrot C. The Multifaceted Role of Protein Phosphatase 1 in Plasmodium. Trends Parasitol, (2020).

25. Kwon YG, Lee SY, Choi Y, Greengard P, Nairn AC. Cell cycle-dependent phosphorylation of mammalian protein phosphatase 1 by cdc2 kinase. Proc Natl Acad Sci U S A 94, 2168–2173 (1997).

26. Bhattacharyya MK, Hong Z, Kongkasuriyachai D, Kumar N. Plasmodium falciparum protein phosphatase type 1 functionally complements a glc7 mutant in Saccharomyces cerevisiae. Int J Parasitol 32, 739–747 (2002).

27. Zhang M, et al. Uncovering the essential genes of the human malaria parasite Plasmodium falciparum by saturation mutagenesis. Science 360, (2018).

28. Bushell E, et al. Functional Profiling of a Plasmodium Genome Reveals an Abundance of Essential Genes. Cell 170, 260–272 e268 (2017).

29. Yokoyama D, Saito-Ito A, Asao N, Tanabe K, Yamamoto M, Matsumura T. Modulation of the growth of Plasmodium falciparum in vitro by protein serine/threonine phosphatase inhibitors. Biochem Biophys Res Commun 247, 18–23 (1998).

30. Paul AS, et al. Co-option of Plasmodium falciparum PP1 for egress from host erythrocytes. Nat Commun 11, 3532 (2020).

31. Fardilha M, Esteves SL, Korrodi-Gregorio L, da Cruz e Silva OA, da Cruz e Silva FF. The physiological relevance of protein phosphatase 1 and its interacting proteins to health and disease. Curr Med Chem 17, 3996–4017 (2010).

32. Hollin T, et al. Essential role of GEXP15, a specific Protein Phosphatase type 1 partner, in Plasmodium berghei in asexual erythrocytic proliferation and transmission. PLoS Pathog 15, e1007973 (2019).

33. Lenne A, et al. Characterization of a Protein Phosphatase Type-1 and a Kinase Anchoring Protein in Plasmodium falciparum. Front Microbiol 9, 2617 (2018).

34. Freville A, et al. Plasmodium falciparum encodes a conserved active inhibitor-2 for Protein Phosphatase type 1: perspectives for novel anti-plasmodial therapy. BMC Biol 11, 80 (2013).

35. Freville A, et al. Identification of a Plasmodium falciparum inhibitor-2 motif involved in the binding and regulation activity of protein phosphatase type 1. FEBS J 281, 4519–4534 (2014).

36. Freville A, et al. Plasmodium falciparum inhibitor-3 homolog increases protein phosphatase type 1 activity and is essential for parasitic survival. J Biol Chem 287, 1306–1321 (2012).

37. Pierrot C, Zhang X, Zanghi G, Freville A, Rebollo A, Khalife J. Peptides derived from Plasmodium falciparum leucine-rich repeat 1 bind to serine/threonine phosphatase type 1 and inhibit parasite growth in vitro. Drug Des Devel Ther 12, 85–88 (2018).

38. Gerald N, Mahajan B, Kumar S. Mitosis in the human malaria parasite Plasmodium falciparum. Eukaryot Cell 10, 474–482 (2011).

39. Guttery DS, Roques M, Holder AA, Tewari R. Commit and Transmit: Molecular Players in Plasmodium Sexual Development and Zygote Differentiation. Trends Parasitol 31, 676–685 (2015).

40. Matthews H, Duffy CW, Merrick CJ. Checks and balances? DNA replication and the cell cycle in Plasmodium. Parasit Vectors 11, 216 (2018).

41. Zeeshan M, et al. Real-time dynamics of Plasmodium NDC80 reveals unusual modes of chromosome segregation during parasite proliferation. J Cell Sci 134, (2020).

42. Arnot DE, Ronander E, Bengtsson DC. The progression of the intra-erythrocytic cell cycle of Plasmodium falciparum and the role of the centriolar plaques in asynchronous mitotic division during schizogony. Int J Parasitol 41, 71–80 (2011).

43. Zeeshan M, et al. Plasmodium kinesin-8X associates with mitotic spindles and is essential for oocyst development during parasite proliferation and transmission. PLoS Pathog 15, e1008048 (2019).

44. Vaughan JA. Population dynamics of Plasmodium sporogony. Trends Parasitol 23, 63–70 (2007).

45. Pandey R, et al. Plasmodium Condensin Core Subunits SMC2/SMC4 Mediate Atypical Mitosis and Are Essential for Parasite Proliferation and Transmission. Cell Rep 30, 1883–1897 e1886 (2020).

46. Wall RJ, et al. Plasmodium APC3 mediates chromosome condensation and cytokinesis during atypical mitosis in male gametogenesis. Sci Rep 8, 5610 (2018).

47. Janse CJ, et al. High efficiency transfection of Plasmodium berghei facilitates novel selection procedures. Molecular and biochemical parasitology 145, 60–70 (2006).

48. Hollin T, De Witte C, Lenne A, Pierrot C, Khalife J. Analysis of the interactome of the Ser/Thr Protein Phosphatase type 1 in Plasmodium falciparum. BMC Genomics 17, 246 (2016).

49. Straschil U, et al. The Armadillo repeat protein PF16 is essential for flagellar structure and function in Plasmodium male gametes. PloS one 5, e12901 (2010).

50. Gelens L, Qian J, Bollen M, Saurin AT. The Importance of Kinase-Phosphatase Integration: Lessons from Mitosis. Trends Cell Biol 28, 6–21 (2018).

51. Nasa I, Rusin SF, Kettenbach AN, Moorhead GB. Aurora B opposes PP1 function in mitosis by phosphorylating the conserved PP1-binding RVxF motif in PP1 regulatory proteins. Sci Signal 11, (2018).

52. Tewari R, et al. The Systematic Functional Analysis of Plasmodium Protein Kinases Identifies Essential Regulators of Mosquito Transmission. Cell Host & Microbe 8, 377–387 (2010).

53. Ceulemans H, Bollen M. Functional diversity of protein phosphatase-1, a cellular economizer and reset button. Physiol Rev 84, 1–39 (2004).

54. Lesage B, Qian J, Bollen M. Spindle checkpoint silencing: PP1 tips the balance. Curr Biol 21, R898–903 (2011).

55. Peggie MW, MacKelvie SH, Bloecher A, Knatko EV, Tatchell K, Stark MJ. Essential functions of Sds22p in chromosome stability and nuclear localization of PP1. J Cell Sci 115, 195–206 (2002).

56. Rodrigues NT, Lekomtsev S, Jananji S, Kriston-Vizi J, Hickson GR, Baum B. Kinetochore-localized PP1-Sds22 couples chromosome segregation to polar relaxation. Nature 524, 489–492 (2015).

57. Verbinnen I, Ferreira M, Bollen M. Biogenesis and activity regulation of protein phosphatase 1. Biochem Soc Trans 45, 89–99 (2017).

58. Sinden RE. Mitosis and meiosis in malarial parasites. Acta Leidensia 60, 19–27 (1991).

59. Solyakov L, et al. Global kinomic and phospho-proteomic analyses of the human malaria parasite Plasmodium falciparum. Nat Commun 2, 565 (2011).

60. Balestra AC, et al. A divergent cyclin/cyclin-dependent kinase complex controls the atypical replication of a malaria parasite during gametogony and transmission. Elife 9, (2020).

61. Invergo BM, Brochet M, Yu L, Choudhary J, Beltrao P, Billker O. Sub-minute Phosphoregulation of Cell Cycle Systems during Plasmodium Gamete Formation. Cell Rep 21, 2017–2029 (2017).

62. Moreira S, Osswald M, Ventura G, Goncalves M, Sunkel CE, Morais-de-Sa E. PP1-Mediated Dephosphorylation of Lgl Controls Apical-basal Polarity. Cell Rep 26, 293–301 e297 (2019).

63. Dessens JT, et al. CTRP is essential for mosquito infection by malaria ookinetes. EMBO J 18, 6221–6227 (1999).

64. Dessens JT, et al. SOAP, a novel malaria ookinete protein involved in mosquito midgut invasion and oocyst development. Molecular microbiology 49, 319–329 (2003).

65. Poulin B, et al. Unique apicomplexan IMC sub-compartment proteins are early markers for apical polarity in the malaria parasite. Biol Open 2, 1160–1170 (2013).

66. Wall RJ, et al. SAS6-like protein in Plasmodium indicates that conoid-associated apical complex proteins persist in invasive stages within the mosquito vector. Sci Rep 6, 28604 (2016).

67. Philip N, Waters AP. Conditional Degradation of Plasmodium Calcineurin Reveals Functions in Parasite Colonization of both Host and Vector. Cell Host Microbe 18, 122–131 (2015).

68. Sebastian S, et al. A Plasmodium calcium-dependent protein kinase controls zygote development and transmission by translationally activating repressed mRNAs. Cell Host Microbe 12, 9–19 (2012).

69. Beetsma AL, van de Wiel TJ, Sauerwein RW, Eling WM. Plasmodium berghei ANKA: purification of large numbers of infectious gametocytes. Exp Parasitol 88, 69–72 (1998).

70. Guttery DS, et al. A putative homologue of CDC20/CDH1 in the malaria parasite is essential for male gamete development. PLoS Pathog 8, e1002554 (2012).

71. Ferguson DJ, et al. Maternal inheritance and stage-specific variation of the apicoplast in Toxoplasma gondii during development in the intermediate and definitive host. Eukaryot Cell 4, 814–826 (2005).

72. Roques M, et al. Plasmodium centrin PbCEN-4 localizes to the putative MTOC and is dispensable for malaria parasite proliferation. Biol Open 8, (2019).

73. Bolger AM, Lohse M, Usadel B. Trimmomatic: a flexible trimmer for Illumina sequence data. Bioinformatics 30, 2114–2120 (2014).

74. Kim D, Langmead B, Salzberg SL. HISAT: a fast spliced aligner with low memory requirements. Nature methods 12, 357–360 (2015).

75. Liao Y, Smyth GK, Shi W. featureCounts: an efficient general purpose program for assigning sequence reads to genomic features. Bioinformatics 30, 923–930 (2014).

76. Robinson MD, McCarthy DJ, Smyth GK. edgeR: a Bioconductor package for differential expression analysis of digital gene expression data. Bioinformatics 26, 139–140 (2010).

77. Ritchie ME, et al. limma powers differential expression analyses for RNA-sequencing and microarray studies. Nucleic acids research 43, e47 (2015).

78. Love MI, Huber W, Anders S. Moderated estimation of fold change and dispersion for RNA-seq data with DESeq2. Genome Biol 15, 550 (2014).

