## Supplementary figures and images for "Protein Phosphatase 1 regulates atypical mitotic and meiotic division in *Plasmodium* sexual stages"

### FigS1

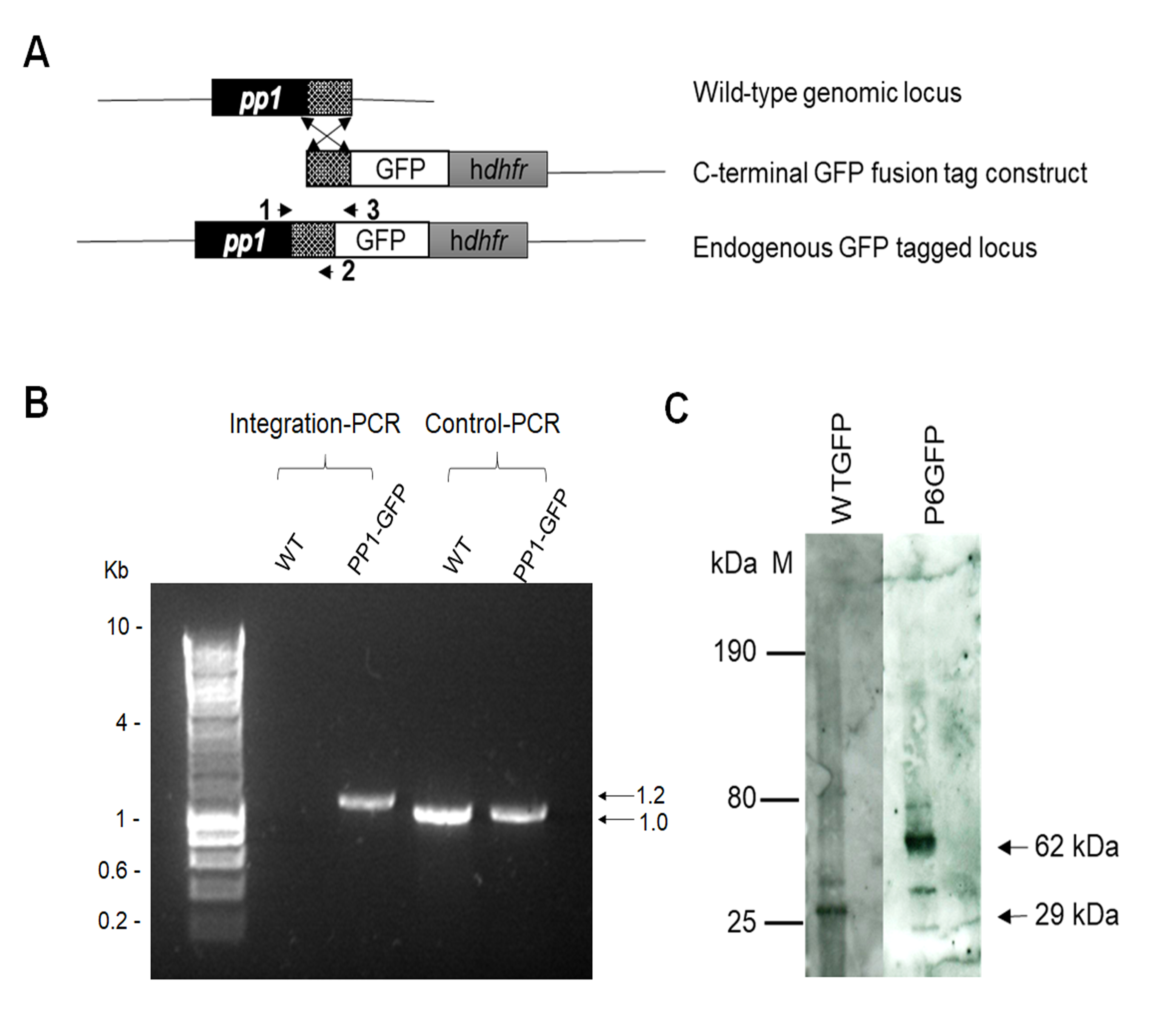

### FigS2

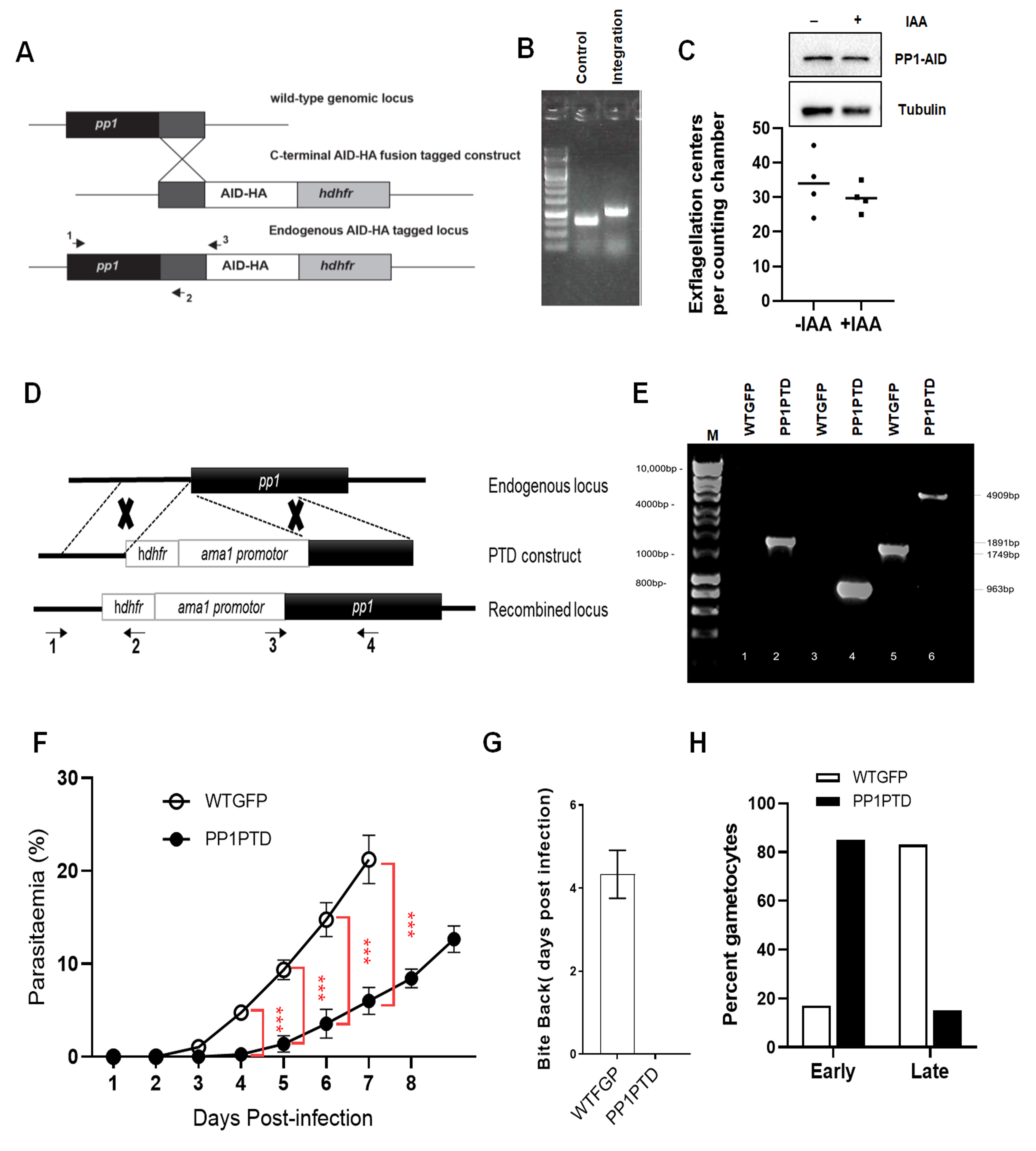

### FigS3

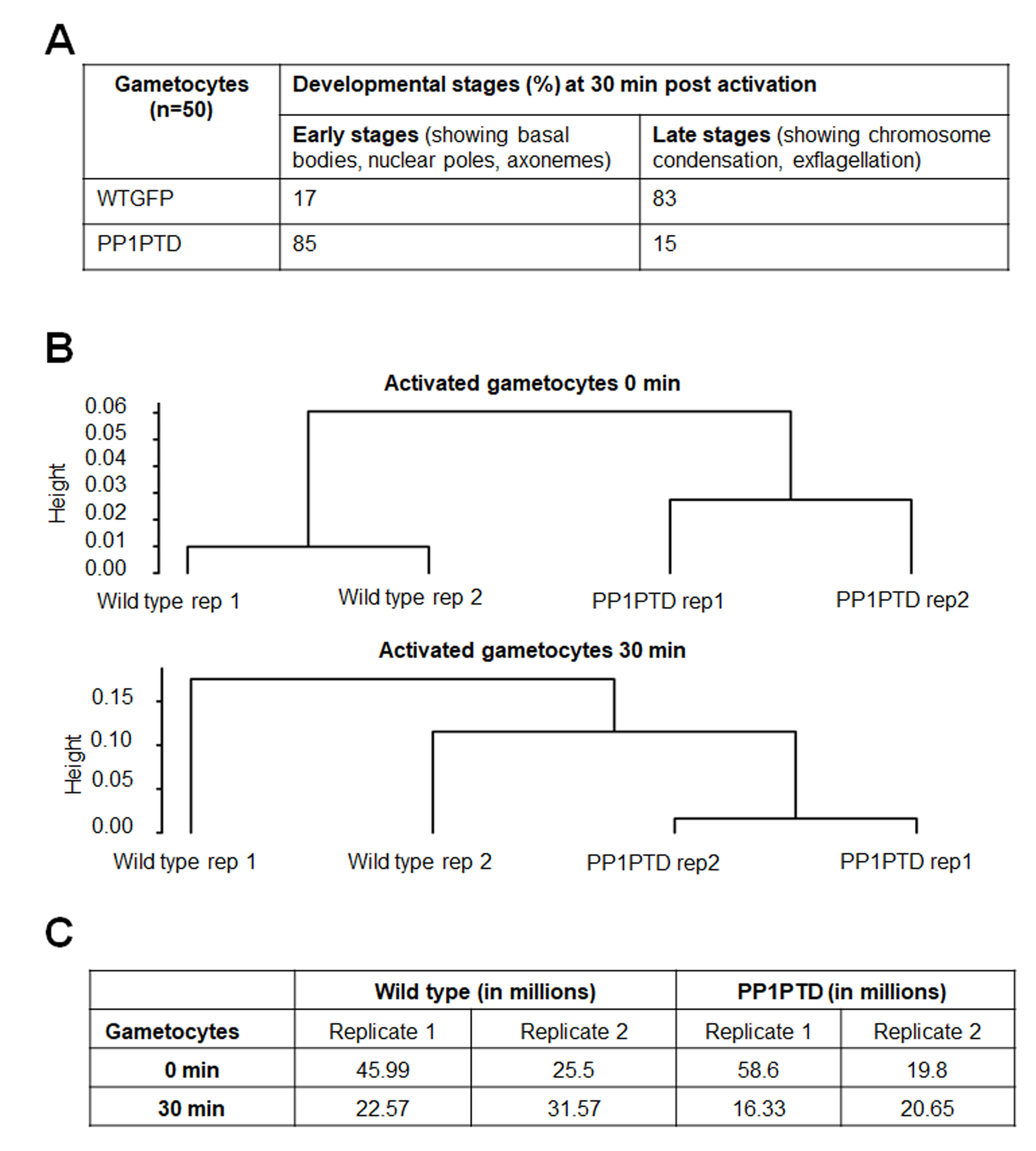
